# Wide distribution of Mediterranean and African Spotted fever agents and the first identification of Israeli Spotted Fever agent in ticks in Uganda

**DOI:** 10.1101/2023.03.29.534855

**Authors:** Wilfred Eneku, Bernard Erima, Anatoli Maranda Byaruhanga, Gladys Atim, Titus Tugume, Qouilazoni A. Ukuli, Hannah Kibuuka, Edison Mworozi, Christina Burrows, Jeffrey W. Koehler, Nora G. Cleary, Michael E. von Fricken, Robert Tweyongyere, Fred Wabwire-Mangen, Denis K. Byarugaba

## Abstract

*Rickettsia* microorganisms are causative agents of several neglected emerging infectious diseases transmitted to humans by ticks among other arthropod vectors. In this study, ticks were collected from four geographical regions of Uganda, pooled in sizes of 1-179 ticks based on location, tick species, life stage, host, and time of collection, and were tested by real time PCR for *Rickettsia* species harboured. The tick pools were tested with primers targeting *gltA, 17kDa* and *omp*A genes, followed by Sanger sequencing of *17kDa* and *ompA* genes. Of the 471 tick pools tested, 116 (24.6%) were positive for *Rickettsia* spp. by the *gltA* primers. The prevalence of *Rickettsia* varied by district with Gulu recording the highest (30.1%) followed by Luwero (28.1%) and Kasese had the lowest (14%). Tick pools with highest positivity rates were from livestock (cattle, goats, sheep, and pigs), 26.9%, followed by vegetation 23.1% and pets (dogs and cats) 19.7%. Of 116 *gltA*-positive tick pools, 86 pools were positive using *17kDa* primers of which 48 purified PCR products were successfully sequenced. The predominant *Rickettsia* spp. identified was *R. africae* (n=15) in four tick species, followed by *R. conorii* (n=5) in three tick species (*Haemaphysalis elliptica, Rhipicephalus appendiculatus*, and *Rh. decoloratus*). *Rickettsia conorii* subsp. *israelensis* was detected in one tick pool. These findings indicate that multiple *Rickettsia* spp. capable of causing human illness are circulating in the four diverse geographical regions of Uganda including new strains previously known to occur in the Mediterranean region. Physicians should be informed about *Rickettsia* spp. infections as potential causes for acute febrile illnesses in these regions. Continued and expanded surveillance is essential to further identify and locate potential hotspots with *Rickettsia* spp. of concern.

**Author Summary:** Tick-borne rickettsioses are emerging infectious diseases of public health importance worldwide. Spotted fever rickettsioses transmitted by ticks can cause mild to severe human illness depending on the *Rickettsia* spp. and co-morbidities. Their diagnosis is challenging due to non-specific symptoms particularly in limited resource settings. Little is known about their prevalence in Uganda. Using entomological and molecular tools, we surveyed and studied tick-borne spotted fever rickettsioses in five districts from four diverse eco-regions of Uganda. Overall, 24.6% (116/471) tick pools were positive for *Rickettsia* species. By sequencing the *17kDa* and *ompA* genes of *Rickettsia*, we identified *R. africae* as the most common agent, followed by *R. conorii* and *R. conorii* subsp. *israelensis*. The findings indicate multiple *Rickettsia* spp. that can cause febrile illness in humans are circulating in the four geographically diverse regions of Uganda. Physicians should be aware these agents are potential causes of febrile illness in these areas, particularly in individuals who encounter livestock or their grazing areas.

## Introduction

*Rickettsia* bacteria, transmitted to humans by arthropod vectors, are responsible for multiple emerging infectious diseases globally (1-3). Clinical presentations of *Rickettsia* spp. differ between two groups, the Spotted Fever Group (SFG) transmitted by ticks and the Typhus Group (TG) transmitted by fleas and lice. Diseases caused by *Rickettsia* spp. have varying seroprevalences rates of 8—10% in the east African region and <1—37% worldwide (4-8). These pathogens are increasingly associated with undifferentiated febrile illnesses in humans. Once infection occurs, the bacteria propagates intracellularly in tissue cells, potentially resulting in severe illness and/or death (9-10). Several cases of SFG rickettsioses have been reported in international travellers returning to their home countries, particularly from sub-Saharan Africa and southeast Asia, where many species of *Rickettsia* are endemic (11-14). Historically, these diseases have been poorly studied in sub-Saharan Africa where the largest burden of disease exists, particularly in indigenous populations (15).

Rickettsioses manifest with non-specific signs such as fever, severe headache, skin rash and general malaise, which can often be misdiagnosed as other febrile illnesses or viral diseases. A confirmed diagnosis requires serological or molecular tests to detect antibodies, a lengthy process of the immune response, potentially leading to a false negative test result if testing occurs too early in the bacterial infection (12,16-17). Moreover, equipment and reagents required for testing are expensive and rely on skilled laboratory technicians. Lengthy testing time, technical equipment, and expertise complicate the ability of countries with constrained resources to properly test for *Rickettsia* infections. Cases of febrile illnesses are often over diagnosed as malaria and later proven otherwise by more sensitive and specific PCR assays (18). Distinguishing *Rickettsia* from other pathogenic agents early in the infection allows for timely treatment and informs any necessary public health measures.

Uganda is home to multiple medically relevant arthropod vectors including ticks, fleas, and mites. Diverse species of ticks have been recovered from both animals and the environment in two regions of the country, the northeast and southwest, often carrying medically relevant human and animal pathogens (19-20). We recently reported the abundance and distribution of seven tick species of the *Rhipicephalus, Haemaphysalis, and Amblyomma* genera in the Ugandan cattle corridor with rickettsial pathogens detected (21). Tick-borne *Rickettsia* spp. of human relevance are prevalent in Uganda including *R. africae, R. conorii*, and *R. massiliae* (22-24).

*Rickettsia africae*, the cause of African tick-bite fever (ATBF), is the most common tick-borne bacterial zoonosis reported in travelers returning from sub-Saharan Africa transmitted predominantly by *Amblyomma* spp.(6,11). On the other hand, *R. conorii* mainly transmitted by *Rhipicephalus sanguineus* causes Mediterranean spotted fever (MSF) largely in the endemic Mediterranean region (10). Israeli spotted fever (ISF) is a similar disease to MSF but causes more severe illness is caused by *R. conorii* subsp. *israelensis* and also transmitted by *Rh. sanguineus* (sensu lato) (22). Although these diseases are generally mild and manifest with the common characteristics of rickettsioses, infection often results in hospitalizations and delays the diagnosis of potentially co-infected febrile illnesses (6,22).

Tick population densities and territories of several organisms have changed significantly in the past decade largely due to anthropogenic and environmental changes resulting from climate change (10, 25-26). An increase in population densities of medically important vectors and pathogens is commonly associated with emergence of disease in humans, representing a major public health concern. This also poses a risk to the 58% of Ugandans who derive their livelihoods through livestock keeping, predominantly kept on open grazing (27). Limited knowledge about the ticks associated with SFG rickettsia transmission, their frequencies, and geographical range in Uganda creates challenges in designing appropriate control measures. While there is evidence of widespread *Rickettsia* spp. present throughout sub-Saharan Africa, there is limited data about the type and frequency in Ugandan ticks (21). Therefore, it is essential to characterise ticks and their associated *Rickettsia* spp. to better inform control strategies and contribute to our understanding of tick-borne diseases in Uganda.

## Materials and Methods

### Study sites

Ticks were collected in five districts [Jinja (Eastern Uganda), Kampala (Capital of Uganda), Kasese (Western Uganda), Gulu (Northern Uganda) and Luwero (Central Uganda)] (Fig 1). The selected districts are considered major economic hubs in their respective regions and are geographically and culturally diverse, with high levels of economic heterogeneity. The source of the ticks were livestock (cattle, goats, sheep, pigs), companion animals (dogs and cats), chicken, and from homestead grass environment between April 2017 and September 2018. Ticks on animals or flagged from vegetation were picked with forceps and preserved in 70% ethanol.

**Figure 1:**
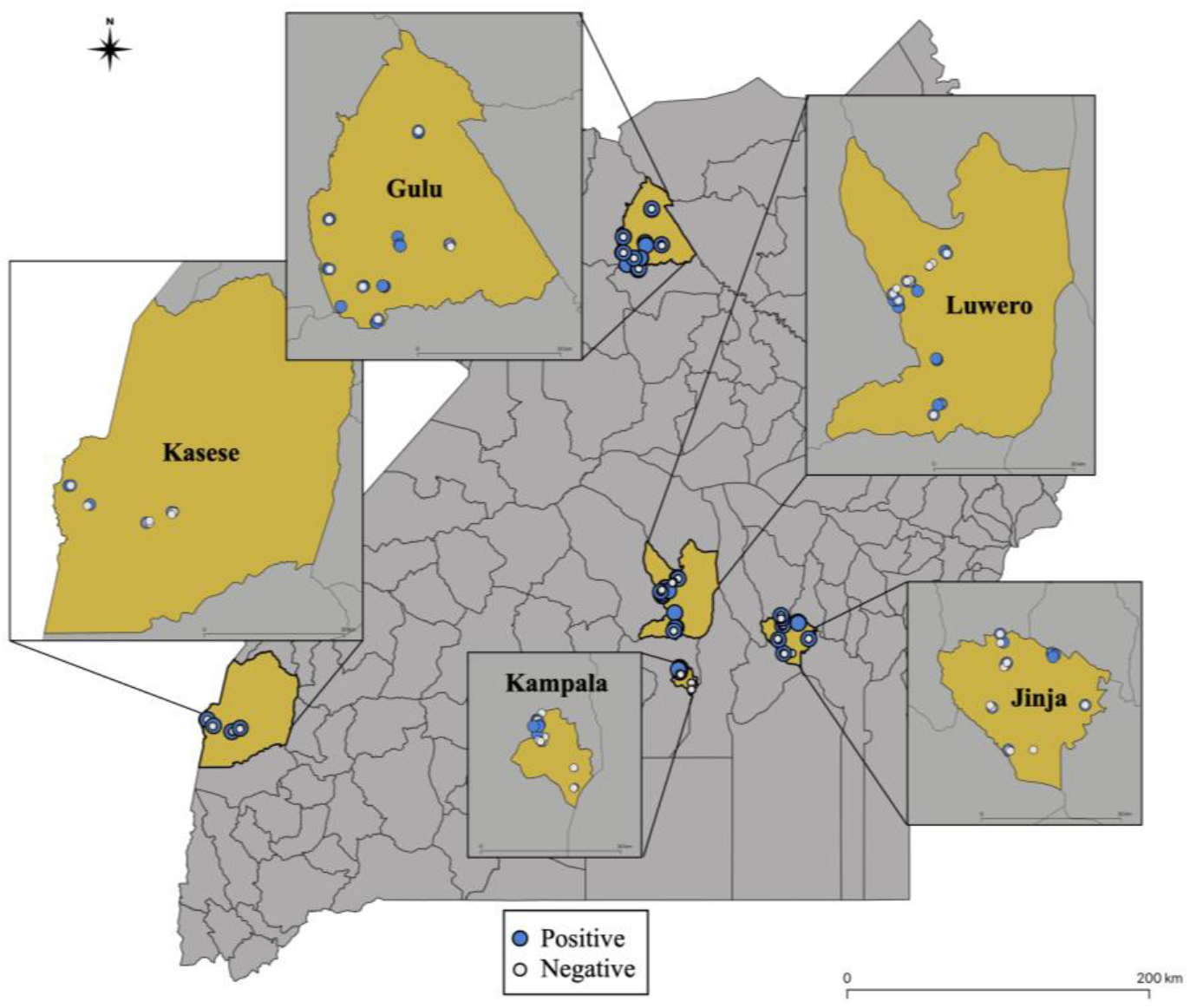
A map of Uganda indicating sites from where pools of ticks were collected and tested for Rickettsia spp. Those tested negative (white circle) and positive (blue circle) are indicated.

### Tick Pools

Collected ticks were identified to the species level using morphologic taxonomic keys (28) under a stereomicroscope. A total of 5,790 ticks were sorted into 471 pools (1-179 ticks per pool, average 12.3) according to tick species, host, collection area, date of collection and developmental stage. There were 306 pools (5,408 ticks) from livestock, 138 pools (677 ticks) from companion animals, 1 pool (1 tick) from a chicken, and 26 pools (64 ticks) from vegetation. The pools were placed in Eppendorf tubes containing RNA later^™^ (Sigma Life Science, Darmstadt, Germany) and disrupted using sterile disposable pestles attached to a motorized grinder (HLD-12, Ryobi, China). Ticks were then homogenized by passing them through 20-gauge needles, with homogenate then stored at -80°C until extracted.

### DNA Extraction and PCR

Total DNA was extracted from the tick homogenates using the Qiagen DNeasy Blood and Tissue kit (Qiagen, Hilden, Germany), according to the manufacturer’s protocol. All 471 tick pool DNA samples were screened for SFG *Rickettsia* spp. with primers amplifying the 74-bp citrate synthase (*gltA*) gene (29). The primers were CS-F (5-TCGCAAATGTTCACGGTACTTT-3) and CS-R (5-TCGTGCATTTCTTTCCATTGTG-3). A second *Rickettsia* genus-specific qPCR amplified a 115-bp segment of the *17kDa* and *ompA* genes to confirm the initial PCR results. *Rickettsia conorii* DNA (provided by Walter Reed Army Institute of Research, Silver Spring, MD) was used as a positive control and ultrapure water as a negative control.

### Sequencing and Phylogenetic Analysis

PCR products were purified using the QIAquick PCR Purification Kit (Qiagen) according to the manufacturer’s instructions. Agarose gel purified amplicons of the *17kDa* and *ompA* genes were sequenced on the SeqStudio (Thermo Fisher Scientific) according to the manufacturer’s recommendations. Forward and reverse reads were aligned using CLC Genomics Workbench (Qiagen) and a consensus sequence for each gene was generated for BLAST analysis. Sequences of *17kDa* and *ompA* genes and references from GenBank were imported and aligned in Geneious Prime 2022.11.0.14.1. The sequences were MAFFT aligned and exported to MEGA 10.2.6 (30) where maximum likelihood trees were created at 1,000 bootstrap iterations.

### Mapping

Descriptive maps showing the collection sites were created in QGIS 3.28 (31). The Uganda district shapefile is available at https://data.unhcr.org/en/documents/details/83043.

### Statistical analysis

The probability of *Rickettsia* spp. detection from the pooled tick samples was estimated using detection rates, maximum likelihood estimation (MLE) and minimum infection rate (MIR) by collection district and tick species. Both MLE and MIR estimates and their corresponding confidence intervals were calculated accounting for individual pool sample sizes in Excel using the CDC’s Mosquito Surveillance Software (https://www.cdc.gov/westnile/resourcepages/mosqSurvSoft.html). A Pearson chi-squared test was used to detect any differences between the distributions of outcomes in different groups, with a p-value of <0.05 considered significant. Data were analyzed using STATA software, version 16.1 (StataCorp, College Station, TX).

## Results

### Distribution of tick species by collection sites

Five tick species were identified from the collections across the five districts. *Rhipicephalus* genera ticks accounted for over half of collections from each district. The most abundant tick species was *Rh. appendiculatus* which constituted 30.6% of all tick pools collected, followed by *Rh. decoloratus* (28.2%) and the least collected tick by pool count was *Rh. sanguineus* (0.8%). *Rhipicephalus sanguineus* was collected from dogs in three districts (Gulu, Jinja, and Kampala) whereas the other four species (*Rh. appendiculatus, Rh. decoloratus, A. variegatum* and *H. elliptica*) were found on animals and environment in all the districts (Figure 2). The tick species variation per district was significant (*X*^2^=32.88, df=20, p= 0.035). Seventy-nine tick pools could not be fully identified because they contained larvae and nymphs with incomplete body parts.

**Figure 2:**
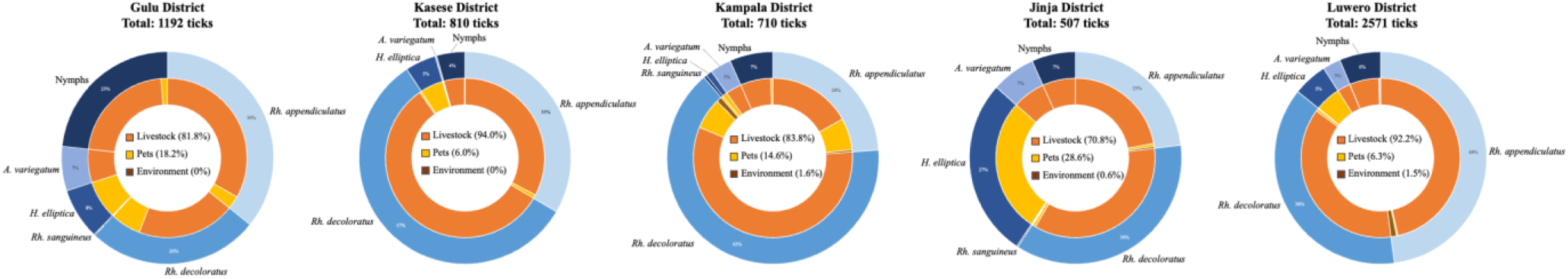
Distribution of ticks by collection district and host. The total number of ticks collected in each district is listed below the respective district name. The outer circle represents the percentage of each tick species from the respective district. The inner circle represents the host distribution of each respective tick species. The total host distribution by district is shown by the percentages in the middle. Livestock includes the chicken recorded from Gulu district. Labels for percentages less than or equal to 1 were excluded.

### Prevalence of *Rickettsia* spp. in the tick pools

Calculated Maximum likelihood estimates (MLE) and Minimum infection rates (MIR) of *Rickettsia* spp. can be found in Table 1. The overall pool positivity rate was 24.6% (116/471) for all the districts, with detection rates varying by site and host type. Livestock (cattle, goats, sheep, and pigs) had the highest pool positivity at 26.9% [(95% CI 21.99, 31.90), (n=83/308)], followed by vegetation 23.1% [(95% CI 6.88, 39.27), (n=6/26)] and domestic animals, 19.7% [(95% CI 13.05, 26.37), (n=27/137)]. The one *Rh. decoloratus* tick pool obtained from a chicken was positive. The MLE for *Rickettsia* spp. by district is as follows: Gulu district had the highest MLE of 4.8% (95% CI: 3.6–6.2) with a corresponding MIR of 3.7% (95% CI: 2.6–4.8) while Kasese had the lowest MLE of 1.1% (95% CI: 0.6–2.0) with a MIR of 1.0% (95% CI: 0.3–1.7). In general, higher MLE values were obtained in districts in the northern region. *Rickettsia*-positive tick pools were from three genera, *Amblyomma* (48.8%), *Rhipicephalus* (24.2%), and *Haemaphysalis* (17.1%). *Amblyomma variegatum* had the highest MLE and MIR across tick species while other tick species had variable MLEs based on districts of collection (Table 1).

**Table 1:** Maximum Likelihood Estimates (MLE) and Minimum Infection Rate (MIR) with corresponding 95% confidence intervals for detection rates of *Rickettsia* in all tick pools

| District | Tick species | Positive pools (%) |  | Total ticks | MLE |  |  | MIR |  |  |
| --- | --- | --- | --- | --- | --- | --- | --- | --- | --- | --- |
|  |  |  |  |  | Point | Low | High | Point | Low | High |
| **Gulu** | *A. variegatum* | 9/14 | (64%) | 81 | 23.8 | 12.6 | 36.6 | 11.1 | 4.3 | 18.0 |
|  | *H. elliptica* | 3/26 | (12%) | 93 | 3.5 | 1.2 | 9.3 | 3.2 | 0 | 6.8 |
|  | *Rh. appendiculatus* | 10/43 | (23%) | 427 | 2.8 | 1.5 | 4.6 | 2.3 | 0.9 | 3.8 |
|  | *Rh. decoloratus* | 13/32 | (41%) | 312 | 6.1 | 3.5 | 9.4 | 4.2 | 2.0 | 6.4 |
|  | *Rh. sanguineus* | 0/1 | (0%) | 2 | 0 | 0 | 54.6 | 0 | - | - |
|  | Nymphs | 9/30 | (30%) | 277 | 4.4 | 2.3 | 7.3 | 3.3 | 1.2 | 5.3 |
|  | **Total** | **44/146** | **(30%)** | **1192** | **4.8** | **3.6** | **6.2** | **3.7** | **2.6** | **4.8** |
| **Kasese** | *A. variegatum* | 0/2 | (0%) | 2 | 0 | 0 | 65.8 | 0 | - | - |
|  | *H. elliptica* | 0/9 | (0%) | 38 | 0 | 0 | 7.5 | 0 | - | - |
|  | *Rh. appendiculatus* | 4/22 | (18%) | 271 | 1.6 | 0.6 | 3.3 | 1.5 | 0 | 2.9 |
|  | *Rh. decoloratus* | 4/18 | (22%) | 464 | 1.0 | 0.4 | 2.2 | 0.9 | 0 | 1.7 |
|  | Nymphs | 0/6 | (0%) | 35 | 0 | 0 | 6.0 | 0 | - | - |
|  | **Total** | **8/57** | **(14%)** | **810** | **1.1** | **0.6** | **2.0** | **1.0** | **0.3** | **1.7** |
| **Kampala** | *A. variegatum* | 2/2 | (100%) | 23 | - | - | - | 8.7 | 0 | 20.2 |
|  | *H. elliptica* | 0/3 | (0%) | 7 | 0 | 0 | 28.0 | 0 | - | - |
|  | *Rh. appendiculatus* | 1/20 | (5%) | 169 | 0.6 | 0.1 | 2.9 | 0.6 | 0 | 1.8 |
|  | *Rh. decoloratus* | 7/29 | (24%) | 461 | 1.7 | 0.8 | 3.1 | 1.5 | 0.4 | 2.6 |
|  | *Rh. sanguineus* | 0/2 | (0%) | 3 | 0 | 0 | 49.9 | 0 | - | - |
|  | Nymphs | 3/11 | (17%) | 47 | 6.4 | 2.3 | 14.5 | 6.4 | 0 | 13.4 |
|  | **Total** | **13/67** | **(19%)** | **710** | **2.0** | **1.2** | **3.2** | **1.8** | **0.9** | **2.8** |
| **Jinja** | *A. variegatum* | 4/8 | (50%) | 34 | 16.6 | 6.0 | 32.1 | 11.8 | 0.9 | 22.6 |
|  | *H. elliptica* | 2/11 | (18%) | 138 | 2.3 | 0.5 | 7.7 | 1.5 | 0 | 3.4 |
|  | *Rh. appendiculatus* | 1/15 | (7%) | 117 | 0.9 | 0.2 | 4.2 | 0.9 | 0 | 2.5 |
|  | *Rh. decoloratus* | 5/20 | (25%) | 183 | 3.0 | 1.3 | 6.0 | 2.7 | 0.4 | 5.1 |
|  | *Rh. sanguineus* | 0/1 | (0%) | 1 | 0 | 0 | 79.4 | 0 | - | - |
|  | Nymphs | 0/7 | (0%) | 34 | 0 | 0 | 7.5 | 0 | - | - |
|  | **Total** | **12/62** | **(19%)** | **507** | **2.8** | **1.6** | **4.7** | **2.4** | **1.0** | **3.7** |
| **Luwero** | *A. variegatum* | 5/15 | (21%) | 70 | 7.5 | 3.4 | 13.6 | 7.1 | 1.1 | 13.2 |
|  | *H. elliptica* | 7/21 | (44%) | 135 | 6.9 | 3.2 | 12.8 | 5.2 | 1.5 | 8.9 |
|  | *Rh. appendiculatus* | 15/44 | (34%) | 1232 | 1.6 | 1.0 | 2.3 | 1.2 | 0.6 | 1.8 |
|  | *Rh. decoloratus* | 8/34 | (24%) | 972 | 0.9 | 0.5 | 1.5 | 0.8 | 0.3 | 1.4 |
|  | Nymphs | 4/25 | (16%) | 162 | 2.7 | 1.1 | 6.0 | 2.5 | 0 | 4.9 |
|  | **Total** | **39/139** | **(28%)** | **2571** | **1.8** | **1.4** | **2.3** | **1.5** | **1.0** | **2.0** |
| **All districts** | **Total** | **116/471** | **(25%)** | **5790** | **2.4** | **2.0** | **2.8** | **2.0** | **1.6** | **2.4** |

### *Rickettsia* spp. identified by nucleotide sequences and phylogenetic analyses

Of the 86 tick pools positive from *gltA* and *17kDa*, 48 pools of purified PCR amplicons were successfully sequenced. The nucleotide sequences obtained from *17kDa* and *ompA* were compared to those available on NCBI GenBank database by BLASTn analyses, with sequence identity and phylogenetic trees presented in Table 2 and Figures 3 & 4. Using the *17kDa* and *ompA* genes, five *Rickettsia* spp. were identified from the tick pools; *R. africae, R. conorii, R. conorii* subsp. *israelensis, R. asembonensis*, and *R. helvetica*. The predominant *Rickettsia* spp. identified was *R. africae*, which was detected in four tick species (*A. variegatum, Rh. appendiculatus, Rh. decoloratus* and *H. elliptica)*, followed by *R. conorii* in three tick species (*Rh. appendiculatus, Rh. decoloratus*, and *H. elliptica*). *Rickettsia conorii* subsp. *israelensis* was also identified in one nymph pool containing ticks removed from a cat. *Rickettsia africae* were recovered from all animal types excluding cats, with *R. conorii* detected in ticks removed from cattle, goats, dogs, and a cat. *Rickettsia asembonensis* was found in *Rh. appendiculatus* and *Rh. decoloratus* from livestock, grass, and a dog and *R. helvetica* was detected in *Rh. appendiculatus* collected from the environment.

**Table 2:**
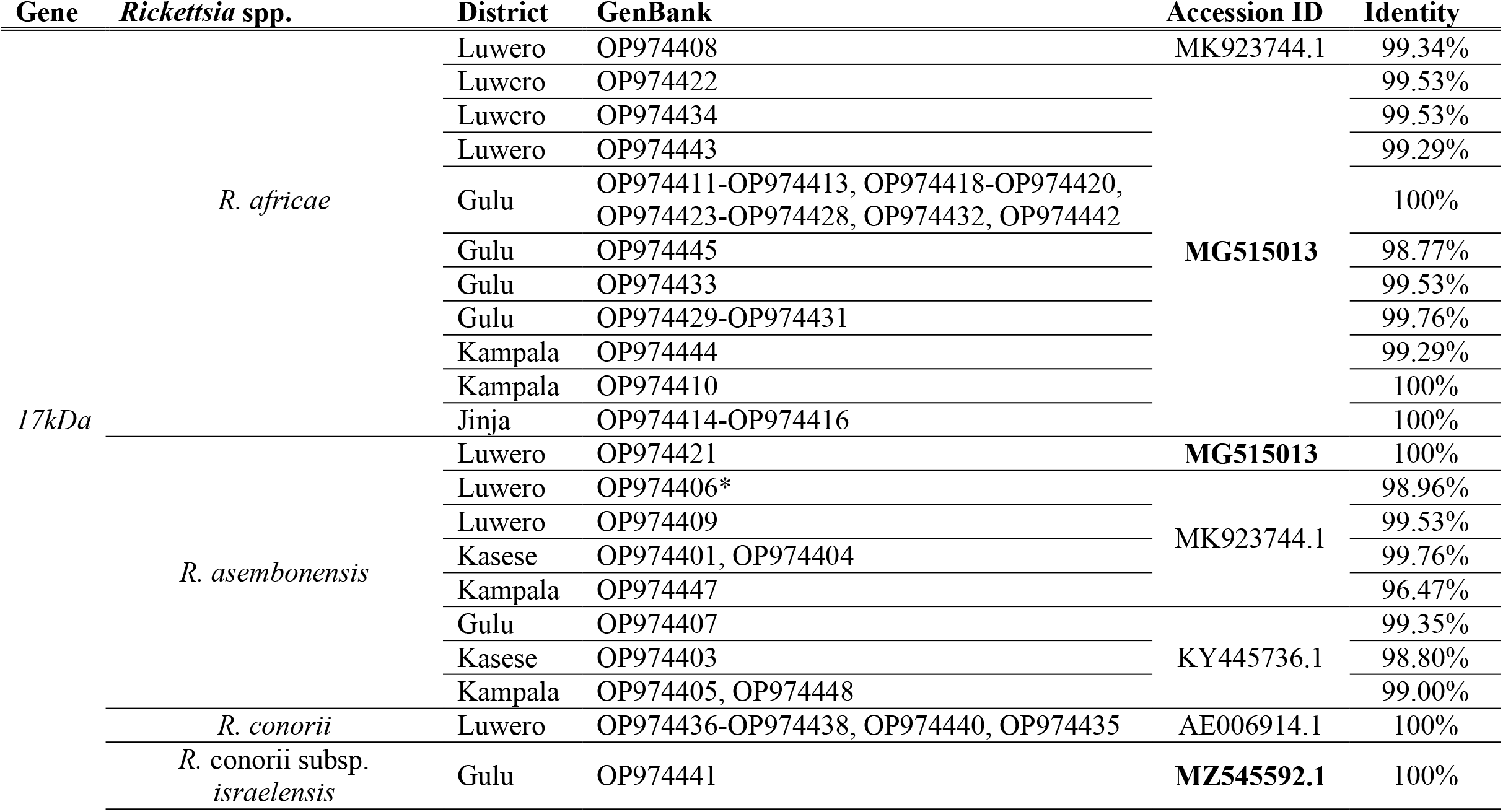
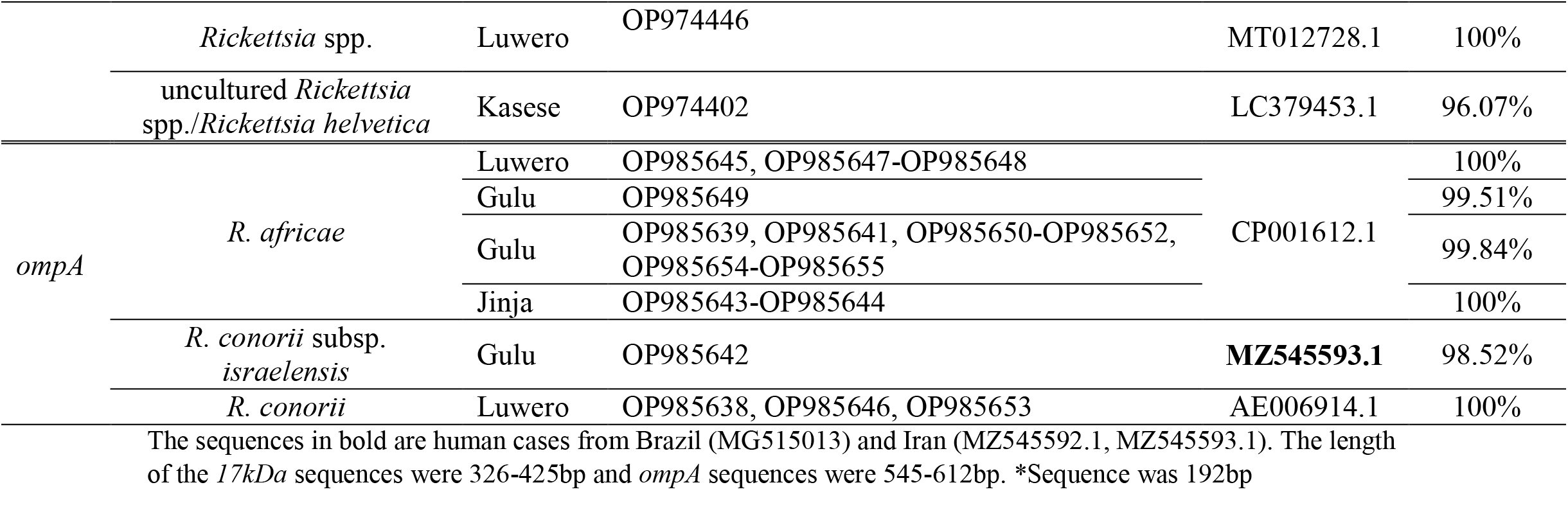
*Rickettsia* spp. identified with the corresponding GenBank accession numbers and identity to sequences on GenBank.

**Figure 3:**
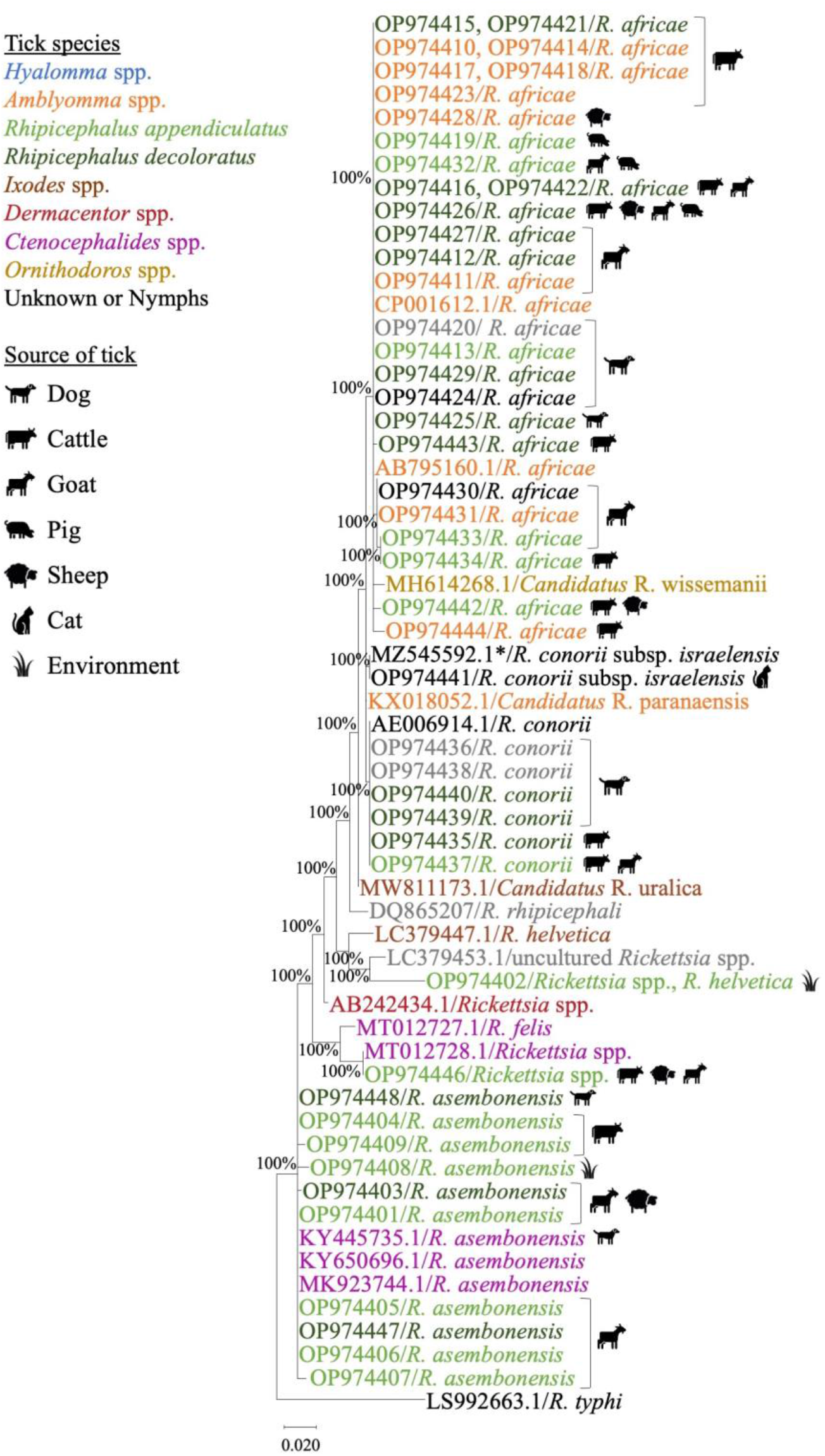
Maximum likelihood tree of the *17kDa Rickettsia* spp. gene using the Tamura-Nei model with sequences ranging from 192 to 426bp. Values less than 70% were excluded from the tree. The legend shows the source of isolation by tick species and tick host by symbol. If multiple source icons appear next to an accession number, the pool of ticks came from more than one source. *One sample, MZ545592.1 was isolated from human serum and not from a tick. All GenBank accession numbers beginning with OP were sequenced in this study.

**Figure 4:**
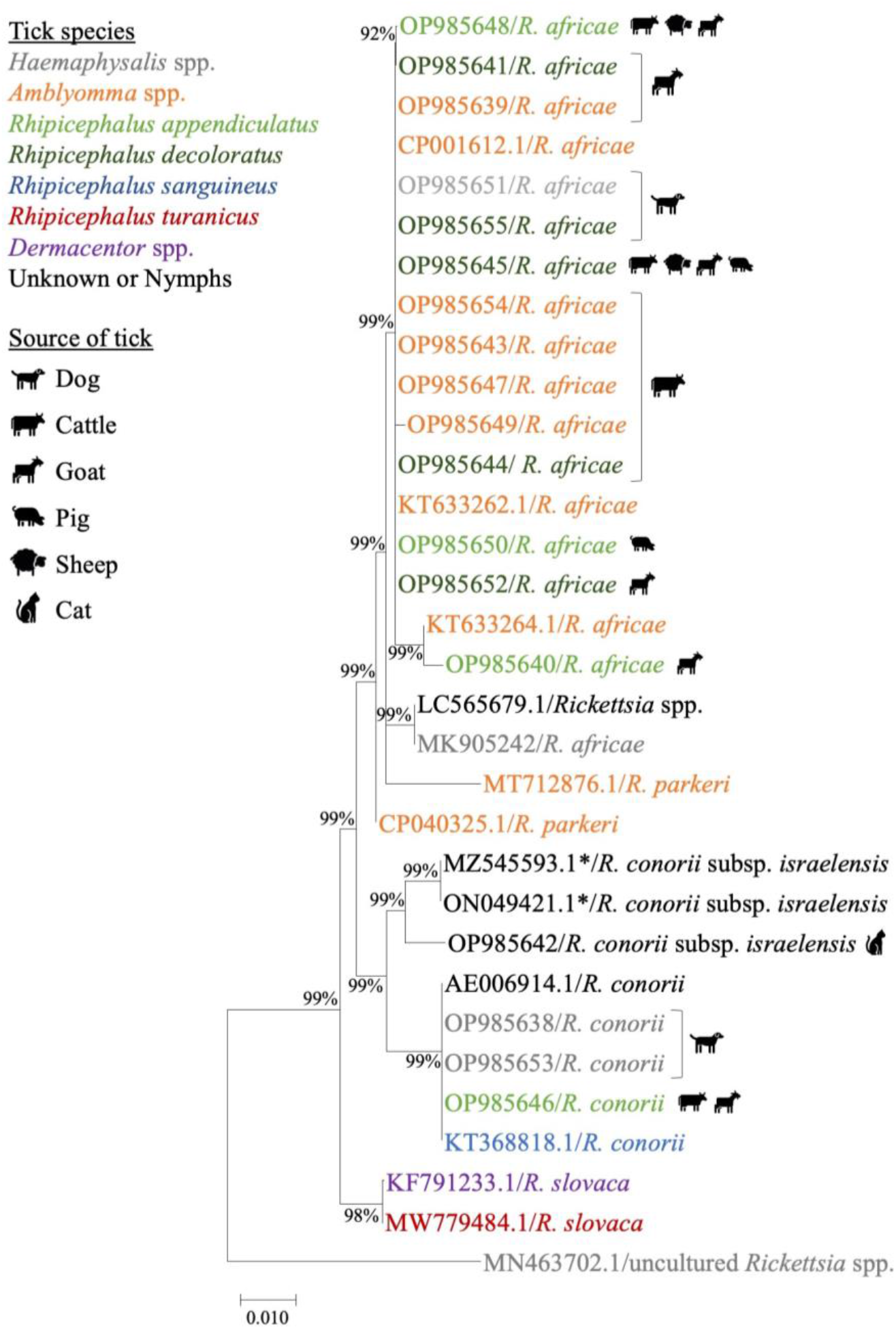
Maximum likelihood tree of the *ompA Rickettsia* spp. gene using the Tamura-Nei model with sequence lengths ranging from 545 to 593bp. Values less than 70% were excluded from the tree. The legend shows the source of isolation by tick species and tick host by symbol. If multiple source icons appear next to an accession number, the pool of ticks came from more than one source. *One sample, MZ545592.1 was isolated from human serum and not from a tick. All GenBank accession numbers beginning with OP were sequenced in this study.

Ticks collected from all livestock species had similar sequences for *R. africae, R. conorii*, and *R. asembonensis* based on the *17kDa* gene. The two sequences obtained from ticks collected from the environment were unique compared to previously published sequences. The comparison of the *17kDa* sequence from this study revealed that tick pools from Gulu, Jinja, Luwero and Kampala had identical homology with *R. africae* strain PELE, which was isolated from a human traveler in Brazil. *Rickettsia conorii* subsp. *israelensis* detected in the Gulu district matched fatal human case detected in Iran. *R. asembonensis 17kDa* sequences obtained from ticks in this study were highly similar to that from flea samples from South America. The *R. helvetica* sequence from this study was unique compared to sequences published on GenBank. The *ompA* sequence comparison revealed *R. africae* from this study were identical to a sequence isolated from *A. variegatum* ticks from Ethiopia and Benin. Additionally, the *R. conorii* sequences were identical to one from a *Rh. sanguineus* tick from a dog in Romania.

## Discussion

A relatively high pool positivity rate (24.6%) for *Rickettsia* spp. was detected in this study. *Rickettsia*-positive ticks were found in every district, with higher MLEs and pool-positivity rates observed in Gulu district (northern Uganda) and Jinja district (eastern Uganda) suggesting potential hotspots for *Rickettsia* spp. infections that need to be further investigated. The Rickettsia spp positivity documented in our study is comparable to a similar study in Kenya that demonstrated 25% *Rickettsia* spp. prevalence in tick pools collected from livestock and camels in dispersed pastoral communities (32). The highest positive pool detection rates among tick species in our study were in *A. variegatum* (48.8%), *Rh. decoloratus* (27.8%) and *Rh. appendiculatus* (21.6%). These ticks feed predominantly on cattle, sheep, goats and large wild ruminants (20). There is evidence of up to 97% prevalence of *R. africae* in *A. variegatum* collected from cattle in Eastern parts of Uganda (29). Similar to other findings from sub-Saharan Africa, our study confirms that *Rickettsia* spp. are likely present across Uganda where animals reside, posing a risk to over half of the Ugandan population that derive their livelihoods from animals (27, 33-35). Further seroprevalence studies would provide more information about the geographic spread of *Rickettsia* pathogens in livestock that could be transmitted to ticks.

The relative variation in infection rates by district could be explained by differences in livestock populations and intensity of acaricide use. Farms in central (Luwero) and western (Kasese) Uganda have more dairy cattle and use more acaricides compared to livestock farms in northern and eastern Uganda, which are predominantly indigenous breeds (36-37). This could also account for the greater tick-species diversity in Gulu as opposed to the other districts. Areas that are using acaracides on their livestock, could be reducing the number and diversity of susceptible tick species. However, tick species that are resistant to acaricides are likely to persist in Uganda including *Amblyomma* spp. and *Rhipicephalus* spp. (38). *Amblyomma* spp. are the major vector for *R. africae* in sub-Saharan Africa. In Gulu district, the highest MLE by tick species was in *A. variegatum* at 23.8 (95% CI: 12.6, 36.6), likely harboring many *Rickettsia* pathogens. Potential resistance in this species may contribute to the maintenance of the highly prevalent *R. africae* in the northern region. *Rhipicephalus* ticks are the most prevalent in Uganda on livestock and were the most common tick genus collected from every district (37). Within this genus, all *Rickettsia* spp. detected in this study were present so they could pose a larger threat if they developed resistance. There have been detections of acaracide resistant *Rhipicephalus* spp. in northern Uganda brought by livestock movements (39). Livestock trading could lead to the movement of ticks, potentially with acaracide resistance, across district or country borders.

While livestock rearing increases the risk for SFG rickettsia, another big industry in Uganda at risk for tick-borne diseases is tourism. Specifically, ATBF has been reported in a Slovenian traveller returning from south western Uganda (14). One major tourism area in the Kasese district, Queen Elizabeth National park, has an estimated 34,000 visitors annually (40). This environment is conducive to wildlife and domestic encounters increasing the risk for human contact with infected ticks. Surprisingly, Kasese district has the lowest MLE at 1.1 (95% CI: 0.6, 2.0), given the large number of tourists visiting the area. Additional sampling should be done in this district to understand the risk of *Rickettsia* spp. to tourists as *R. africae* was not detected in this region, likely because only two *A. variegatum* ticks were collected. Travelers should be aware of the risk of illness associated with these pathogens.

The detection of *Rickettsia* sequences (*R. africae* and *R. conorii*) with high homology to ones causing human illness in Brazil and Iran emphasizes the importance of monitoring these pathogens in Uganda. The *R. africae* strain, PELE, caused ATBF leading to hospitalization, in a traveler from South Africa and was identical to *17kDa* sequences from this study from multiple districts (11). Of concern is the *R. conorii* subsp. *israelensis* sequence from Gulu district that matched a fatal human case from Iran (41). The discovery of these pathogens in the most abundant tick species on livestock (37) poses a high risk to Ugandans as it has the potential to infect humans and cause human illness.

As the first report of *R. africae* in Uganda in *Rh. appendiculatus* and *Rh. decoloratus* in all collection districts aside from Kasese, there is potential for more insight into the distribution of *Rickettsia* spp. among *Rhipicephalus* ticks. Especially because these two *Rhipicephalus* spp. Are the most abundant livestock ticks in Uganda, which was confirmed in this study and the most multi-acaricide resistant ticks on animal farms (20, 38). This study also presents the first detection of *R. conorii* in *Rh. appendiculatus* and *Rh. decoloratus* ticks in Uganda, which could lead to MSF, especially since a common transmission route is contact with domestic animals (22). Another causative agent of febrile illness, *R. helvetica*, was found for the first time in Uganda in *Rh. appendiculatus* and was unique to other published sequences. Interestingly, this tick was collected from vegetation and not from livestock. *Rickettsia asembonensis* was detected in two tick species, *Rh. appendiculatus* and *Rh. decoloratus*, but it is mostly flea borne. It has occasionally been detected in ticks with limited information about its pathogenicity in humans (42). Additional studies on *Rhipicephalus* genera would be beneficial to understanding the scope of these *Rickettsia* spp. in these ubiquitous vectors across Uganda.

### Limitations

Ticks were collected from five districts in Uganda to represent the four regions. Careful consideration was taken when extrapolating the results from the districts as they may not be representative of *Rickettsia* spp. found within the entire respective region. Additionally, a limited number of ticks (64/5790) were collected from the vegetation and analyzed in 26/471 pools so minimal environmental conclusions were made in this study. Ticks were identified solely using morphology thus limiting the confidence of species identification. By pooling ticks, the MLE was immeasurable when 100% of tick pools were positive and MIR was immeasurable when 0% of tick pools were positive and this was noted in Table 2 using dashes (-). The PCR targets were designed for one species, *Rickettsia*, so co-infection was not possible to assess.

## Conclusions

This is the first major surveillance study using targeted gene sequencing for *Rickettsia* spp. covering diverse ecological zones of Uganda. Several *Rickettsia* spp. were detected circulating in multiple tick species in the four regions including the first detection of the ISF agent in ticks in Uganda. Additionally, the ATBF and MSF causative agents were identified in Uganda for the first time in *Rh. appendiculatus and Rh. decoloratus* ticks. Further pathogen surveillance targeting *Rickettsia* spp. will improve the knowledge of the geographic region of infected ticks in Uganda and track future evolution. Clinicians must be informed of circulating *Rickettsia* spp. endemic to Uganda to timely and effectively detect, treat, and prevent human illness. Since *Rickettsia* spp. often cause febrile illness in patients and can be misdiagnosed as malaria, an investment in enhancing diagnostic capabilities in public health facilities would enable accurate early detection. Consideration should be given to revise the Uganda clinical guidelines to redesignate *Rickettsia* spp. management from the underequipped Health Centre IIs to a higher-level healthcare facility with more qualified medical staff and improved diagnostic tests. The detection of *Rickettsia* spp. in every district surveyed in Uganda highlights the need to monitor the threat of rickettsial disease in these regions and develop rapid diagnostic tests. The lack of diagnostic capacity of equipment and expertise for diagnosis in the healthcare facilities in Uganda. The detection of multiple pathogens known to cause human illness in tick pools from this study highlights the need for essential tick-borne pathogen surveillance. Tick-borne pathogen surveillance and seroprevalence studies are essential in Uganda to further characterize the *Rickettsia* spp. which threated Ugandans, travelers, and public health.

## Acknowledgment

We are grateful to all persons who helped in this study particularly, the vector control officers of the five districts who helped in the collection of ticks from animals and vegetation. We also acknowledge the staff of Makerere University Walter Reed Project that work at the Emerging Infectious Disease Laboratory for the support during laboratory tests for rickettsia on the ticks.

## Data availability statement

The sequences from this study are available on GenBank under the accession numbers: OP974401-OP974448, OP985638-OP985655

## Author Contributions

DKB, JWK, HK, EM, RT and WE: Conceived and designed the study. WE, BE, AMB, GA, TT, QAU, NGC, performed the experiments. WE, NGC, MEvF, DKB, analyzed the data. All the authors participated in drafting and writing the manuscript.

## Supporting Information

**S1 Fig 1. Distribution of tick collection events in the five districts where** tick pools were collected and tested for *Rickettsia* spp. Negative pools (white circle) and positive pools (blue circle) are indicated. (TIFF)

**S2 Fig 2. Distribution of ticks by collection district and host**. The total number of ticks collected in each district is listed below the respective district name. The outer circle represents the percentage of each tick species from the respective district. The inner circle represents the host distribution of each respective tick species. The total host distribution by district is shown by the percentages in the middle. Livestock includes the chicken recorded from Gulu district. Labels for percentages less than or equal to 1 were excluded. (TIFF)

**S3 Fig 3. Maximum likelihood tree of the *17kDa Rickettsia* spp. gene using the Tamura-Nei model with sequences ranging from 192 to 426bp**. Values less than 70% were excluded from the tree. The legend shows the source of isolation by tick species and tick host by symbol. If multiple source icons appear next to an accession number, the pool of ticks came from more than one source. *One sample, MZ545592.1 was isolated from human serum and not from a tick. All GenBank accession numbers beginning with OP were sequenced in this study. (TIFF)

**S4 Fig 4. Maximum likelihood tree of the *ompA Rickettsia* spp. gene using the Tamura-Nei model with sequence lengths ranging from 545 to 593bp**. Values less than 70% were excluded from the tree. The legend shows the source of isolation by tick species and tick host by symbol. If multiple source icons appear next to an accession number, the pool of ticks came from more than one source. *One sample, MZ545592.1 was isolated from human serum and not from a tick. All GenBank accession numbers beginning with OP were sequenced in this study. (TIFF)

**S1 Table**. Maximum Likelihood Estimates (MLE) and Minimum Infection Rate (MIR) with corresponding 95% confidence intervals for detection rates of *Rickettsia* in all tick pools. (DOC)

**S2 Table**. *Rickettsia* spp. identified with the corresponding GenBank accession numbers and identity to sequences on GenBank. (DOC)

## References

1. Raoult D, Roux V. Rickettsioses as paradigms of new or emerging infectious diseases. Clin Microbiol Rev. 1997;10(4):694–719.

2. Parola P, Paddock CD, Raoult D. Tick-borne rickettsioses around the world: emerging diseases challenging old concepts. Clin Microbiol Rev [Internet]. 2005 Oct 1 [cited 2018 Oct 6];18(4):719–56. Available from: http://www.ncbi.nlm.nih.gov/pubmed/16223955

3. Walker DH. Rickettsiae and Rickettsial Infections: The Current State of Knowledge. Clin Infect Dis [Internet]. 2007 Jul 15 [cited 2018 Nov 9];45(Supplement_1):S39–44. Available from: http://academic.oup.com/cid/article/45/Supplement_1/S39/357524/Rickettsiae-and-Rickettsial-Infections-The-Current

4. Prabhu M, Nicholson WL, Roche AJ, Kersh GJ, Fitzpatrick KA, Oliver LD, et al. Q Fever, Spotted Fever Group, and Typhus Group Rickettsioses Among Hospitalized Febrile Patients in Northern Tanzania. Oxford Univ Press. 2011;53.

5. Thiga JW, Mutai BK, Richards AL, Waitumbi JN. High Seroprevalence of Antibodies against in Patients with Febrile Illness, Kenya. Emerg Infect Dis. 2015;21(4):688–91.

6. Parola P, Paddock CD, Socolovschi C, Labruna MB, Mediannikov O, Kernif T, et al. Update on Tick-Borne Rickettsioses around the World: a Geographic Approach. Clin Microbiol Rev [Internet]. 2013 [cited 2018 Oct 16]; Available from: http://www.bacterio.cict.fr/qr

7. Mayxay M, Castonguay-Vanier J, Chansamouth V, Dubot-Pérès A, Paris DH, Phetsouvanh R, et al. Causes of non-malarial fever in Laos: A prospective study. Lancet Glob Heal. 2013;1(1):46–54.

8. Abhilash KPP, Jeevan JA, Mitra S, Paul N, Murugan TP, Rangaraj A, et al. Acute Undifferentiated Febrile Illness in Patients Presenting to a Tertiary Care Hospital in South India: Clinical Spectrum and Outcome. J Glob Infect Dis [Internet]. 2016 [cited 2019 Apr 11];8(4):147–54. Available from: http://www.ncbi.nlm.nih.gov/pubmed/27942194

9. Weinberger M, Keysary A, Sandbank J, Zaidenstein R, Itzhaki A, Strenger C, et al. Fatal Rickettsia conorii subsp. israelensis infection, Israel. Emerg Infect Dis. 2008;14(5):821–4.

10. Sekeyová Z, Danchenko M, Filipčík P, Fournier PE. Rickettsial infections of the central nervous system. PLoS Negl Trop Dis. 2019;13(8):1–18.

11. Angerami RN, Krawczak FS, Nieri-Bastos FA, Santos F, Medorima C, Resende MR, et al. First report of African tick-bite fever in a South American traveler. SAGE Open Med Case Reports. 2018;6:2050313X1877530.

12. Ericsson CD, Jensenius M, Fournier P-E, Raoult D. Rickettsioses and the International Traveler. Clin Infect Dis [Internet]. 2004 Nov 15 [cited 2018 Oct 17];39(10):1493–9. Available from: https://academic.oup.com/cid/article-lookup/doi/10.1086/425365

13. Eldin C, Parola P. Update on tick-borne bacterial diseases in Travelers. Curr Infect Dis Rep. 2018;20(17).

14. Bogovic P, Lotric-furlan S, Korva M, Avsic-zupanc T, Capuzzo C, Felipe CB, et al. African Tick-Bite Fever in Traveler Returning to Slovenia from Uganda Polymyxin B Resistance in Carbapenem-Resistant Klebsiella pneumoniae, São Paulo, Brazil. Emerg Infect Dis. 2016;22(10):1848–9.

15. Stewart AG, Stewart AGA. An update on the laboratory diagnosis of rickettsia spp. Infection. Pathogens. 2021;10(10):1–11.

16. Kovácová E, Kazár J. Rickettsial diseases and their serological diagnosis. Clin Lab [Internet]. 2000 [cited 2018 Oct 17];46(5–6):239–45. Available from: http://www.ncbi.nlm.nih.gov/pubmed/10853230

17. Paris DH, Dumler JS. State of the art of diagnosis of rickettsial diseases: The use of blood specimens for diagnosis of scrub typhus, spotted fever group rickettsiosis, and murine typhus. Curr Opin Infect Dis. 2016;29(5):433–9.

18. Ghai RR, Thurber MI, El Bakry A, Chapman CA, Goldberg TL. Multi-method assessment of patients with febrile illness reveals over-diagnosis of malaria in rural Uganda. Malar J [Internet]. 2016 Dec 7 [cited 2018 Oct 15];15(1):460. Available from: http://malariajournal.biomedcentral.com/articles/10.1186/s12936-016-1502-4

19. Kabi F, Masembe C, Muwanika V, Kirunda H, Negrini R. Geographic distribution of non-clinical Theileria parva infection among indigenous cattle populations in contrasting agro-ecological zones of Uganda: Implications for control strategies. Parasites and Vectors. 2014;7(1):1–9.

20. Byaruhanga C, Collins NE, Knobel D, Kabasa W, Oosthuizen MC. Endemic status of tick-borne infections and tick species diversity among transhumant zebu cattle in Karamoja Region, Uganda: Support for control approaches. Vet Parasitol Reg Stud Reports [Internet]. 2015;1–2:21–30. Available from: http://dx.doi.org/10.1016/j.vprsr.2015.11.001

21. Corrigan J, Marion B, English J, Eneku W, Weng JL, Rugg M, et al. Minimal Rickettsial Infection Rates and Distribution of Ticks in Uganda: An Assessment of the Seasonal Effects and Relevance to Tick-Borne Disease Risk in East Africa. J Med Entomol. 2022;(Xx):1–8.

22. Onyiche TE, Labruna MB, Saito TB. Unraveling the epidemiological relationship between ticks and rickettsial infection in Africa. Front Trop Dis. 2022;3(September):1–29.

23. Spernovasilis N, Markaki I, Papadakis M, Mazonakis N, Ierodiakonou D. Mediterranean spotted fever: Current knowledge and recent advances. Trop Med Infect Dis. 2021;6(4).

24. Proboste T, Kalema-Zikusoka G, Altet L, Solano-Gallego L, Fernández De Mera Ig, Chirife AD, et al. Infection and exposure to vector-borne pathogens in rural dogs and their ticks, Uganda. Parasites and Vectors [Internet]. 2015;8(1):1–9. Available from: http://dx.doi.org/10.1186/s13071-015-0919-x

25. Olwoch JM, Reyers B, Engelbrecht FA, Erasmus BFN. Climate change and the tick-borne disease, Theileriosis (East Coast fever) in sub-Saharan Africa. J Arid Environ. 2008;72(2):108–20.

26. Khatchikian CE, Prusinski M, Stone M, Bryon Backenson P, Wang IN, Levy MZ, et al. Geographical and environmental factors driving the increase in the Lyme disease vector Ixodes scapularis. Ecosphere. 2012;3(10):1–18.

27. Waiswa D, Günlü A, Mat B. Development opportunities for livestock and dairy cattle production in Uganda: a Review. Res J Agric For Sci Int Sci Community Assoc [Internet]. 2021;9(1):18–24. Available from: https://www.isca.me

28. Walker AR, Bouattour A, Camicas J., Estrada-peña A, Horak I., Latif A., et al. Ticks of domestic animals in Africa: a guide to identification of species [Internet]. The University of Edinburgh. 2003. 227 p. Available from: http://www.researchgate.net/publication/259641898_Ticks_of_domestic_animals_in_Africa_a_guide_to_identification_of_species/file/5046352d0429878d7f.pdf

29. Nakao R, Qiu Y, Igarashi M, Magona JW, Zhou L, Ito K, et al. High prevalence of spotted fever group rickettsiae in Amblyomma variegatum from Uganda and their identification using sizes of intergenic spacers. Ticks Tick Borne Dis. 2013;4(6):506–12.

30. Kumar S, Stecher G, Li M, Knyaz C, Tamura K. MEGA X: Molecular evolutionary genetics analysis across computing platforms. Mol Biol Evol. 2018;35(6):1547–9.

31. QGIS.org. QGIS Geographic Information System. QGIS Association. [Internet]. 2022 [cited 2023 Feb 25]. Available from: https://www.qgis.org/en/site/

32. Koka H, Sang R, Kutima HL, Musila L, Macaluso K. The detection of spotted fever group rickettsia DNA in tick samples from pastoral communities in Kenya. J Med Entomol. 2017;54(3):774–80.

33. Adjou Moumouni PF, Terkawi MA, Jirapattharasate C, Cao S, Liu M, Nakao R, et al. Molecular detection of spotted fever group rickettsiae in Amblyomma variegatum ticks from Benin. Ticks Tick Borne Dis [Internet]. 2016;7(5):828–33. Available from: http://dx.doi.org/10.1016/j.ttbdis.2016.03.016

34. Tomassone L, De Meneghi D, Adakal H, Rodighiero P, Pressi G, Grego E. Detection of Rickettsia aeschlimannii and Rickettsia africae in ixodid ticks from Burkina Faso and Somali Region of Ethiopia by new real-time PCR assays. Ticks Tick Borne Dis [Internet]. 2016;7(6):1082–8. Available from: http://dx.doi.org/10.1016/j.ttbdis.2016.09.005

35. Nakao R, Qiu Y, Salim B, Hassan SM, Sugimoto C. Molecular Detection of Rickettsia africae in Amblyomma variegatum Collected from Sudan. Vector-Borne Zoonotic Dis. 2015;15(5):323–5.

36. Vudriko P, Okwee-Acai J, Byaruhanga J, Tayebwa DS, Okech SG, Tweyongyere R, et al. Chemical tick control practices in southwestern and northwestern Uganda. Ticks Tick Borne Dis [Internet]. 2018;9(4):945–55. Available from: https://doi.org/10.1016/j.ttbdis.2018.03.009

37. Kasaija PD, Estrada-Peña A, Contreras M, Kirunda H, de la Fuente J. Cattle ticks and tick-borne diseases: a review of Uganda’s situation. Ticks Tick Borne Dis. 2021;12(5).

38. Vudriko P, Okwee-Acai J, Tayebwa DS, Byaruhanga J, Kakooza S, Wampande E, et al. Emergence of multi-acaricide resistant Rhipicephalus ticks and its implication on chemical tick control in Uganda. Parasites and Vectors [Internet]. 2016;9(1). Available from: http://dx.doi.org/10.1186/s13071-015-1278-3

39. Selby R, Bardosh K, Picozzi K, Waiswa C, Welburn SC. Cattle movements and trypanosomes: Restocking efforts and the spread of Trypanosoma brucei rhodesiense sleeping sickness in post-conflict Uganda. Parasites and Vectors. 2013;6(1):1–12.

40. Dunwiddie L, Shaw RT. Balancing Conservation and Development : A Case Study of Economic Efficiency in Queen Elizabeth National Park, Uganda. 2013;

41. Esmaeili S, Latifian M, Khalili M, Farrokhnia M, Stenos J, Shafiei M, et al. Fatal Case of Mediterranean Spotted Fever Associated with Septic Shock, Iran. Emerg Infect Dis. 2022;28(2):485–8.

42. Kocher C, Morrison AC, Leguia M, Loyola S, Castillo RM, Galvez HA, et al. Rickettsial Disease in the Peruvian Amazon Basin. PLoS Negl Trop Dis. 2016;10(7):1–13.

